# Grazing protists foster zebrafish virus transmission

**DOI:** 10.64898/2026.10.15.759486

**Authors:** Katherine A. Deets, Richard T. Thompson, Hannah J. Young, Claire E. Tobin, James A. Gagnon, Nels C. Elde

**Affiliations:** Department of Human Genetics, University of Utah School of Medicine, Salt Lake City, UT, 84112, USA; School of Biological Sciences, University of Utah, Salt Lake City, UT, 84112, USA; Howard Hughes Medical Institute, 4000 Jones Bridge Road, Chevy Chase, MD 20815, USA

## Abstract

Virus transmission is facilitated by diverse means, ranging from particles hitchhiking on dust and other fomites to vector-borne replication. Aquatic ecosystems are rich with interactions between species at various trophic levels, but how these interactions influence virus transmission and infection biology is largely unknown. By mixing zebrafish picornavirus 1 (ZfPV-1) with a panel of freshwater microbial eukaryotes, we found that phagocytic cells promote virus infections in larval zebrafish, even though the virus is unable to replicate in these microbes. Ciliates, which act as crucial concentrators of micronutrients via continuous grazing behavior, stood out as promoters of robust and accelerated virus infections. We found that simply adding the model ciliate *Tetrahymena thermophila* to mixed populations of infected and uninfected zebrafish facilitated chains of experimental ZfPV-1 transmission between fish. This process depends on *Tetrahymena* phagocytosis and is likely due to the ability of the ciliates to deliver a concentrated bolus of viruses when consumed by larval fish. Notably, the pathogenic snakehead rhabdovirus was not transmitted after mixing with *Tetrahymena*, revealing selectivity, potentially through coevolution with ZfPV-1, that is not evident in all RNA fish viruses. We propose that non-host ciliates can act as virus vessels that increase virus transmission between hosts while potentially altering courses of infection by boosting zebrafish immunity. Virus vessels fill a crucial space on the continuum of virus transmission dynamics for viruses lacking the ability to replicate in intermediate species. Given the tools available to manipulate these genetically accessible systems, this paradigm opens new avenues for investigating otherwise hidden features of infection biology with experiments better approximating natural complexity in the wild.

## Introduction

Transmission between hosts is a fundamental requirement for viruses. While some viruses pass directly from one host to another, many others rely on facilitated transmission, which ranges from passive transport along contaminated surfaces to active replication in intermediate vector species^1^. Viruses traveling between hosts also encounter many non-host organisms that have the potential influence transmission through active processes like ingestion and degradation. However, our understanding of how dynamic biological interactions impact virus transmission remains limited by the challenge of modeling these processes in the lab.

Aquatic environments are particularly rich settings to study facilitated virus transmission across trophic levels. Here, viruses shed from infected vertebrate hosts are immersed in diverse communities of phagocytic protists that are continually consuming, concentrating, and redistributing particles^2–4^. These active biological processes have the potential to alter viral persistence and opportunities for host encounter^5–7^, yet their contribution to transmission and potentially even virus evolution remain largely unexplored.

Ciliates are prime phagocytic candidates for influencing virus transmission given their prevalence and the enormous numbers of suspended particles they process each day^8,9^. These single-celled eukaryotes have long been assumed to simply degrade most consumed viruses^10–14^. However, their potential as a key food source for larval fish^15,16^ and known roles in the persistence and dissemination of a variety of vertebrate bacterial pathogens^17–20^ suggests these grazing organisms may also drive the transmission of aquatic viruses.

To ask how ciliates influence aquatic virus transmission between vertebrate hosts, we studied zebrafish (*Danio rerio*), a workhouse species for developmental genetics that are exposed to a diversity of both viruses and ciliates^21^. The recent discovery and semi-domestication of a widespread, endemic picornavirus, zebrafish picornavirus 1 (ZfPV-1)^22,23^, provides a useful experimental tool to understand modes of virus transmission in complex aquatic systems. ZfPV-1 appears to spread through a fecal-oral route of transmission common to many enteroviruses, making it an appealing candidate for modeling facilitated transmission by non-host species.

We surveyed a panel of ciliates and other microbial eukaryotes, including the model ciliate *Tetrahymena thermophila*^24^, and found that ciliate-mediated delivery of ZfPV-1 altered infection kinetics and antiviral responses in larval fish compared to exposures to free virus. This form of facilitated virus transmission depends on intact *Tetrahymena* cells and active phagocytosis, revealing a process where non-host cells consume viral particles from the water and deliver them to susceptible hosts. Furthermore, the presence of *Tetrahymena* among populations of infected and uninfected larval fish increased virus transmission among individuals that would otherwise not sustain chains of transmission. We coin these non-host organisms that play an active role in facilitated transmission as “virus vessels”—a means of spread distinct from vectors that act as an intermediary species for virus.

Modeling the diverse routes by which viruses gain access to new hosts is critical for understanding how infectious diseases move between susceptible populations. By studying virus infections in the presence of the phagocytic ciliate *Tetrahymena*, we discovered a far-reaching role for non-host microbial eukaryotes in facilitated transmission and established a framework for dissecting how dynamic biological interactions influence virus spread.

## Results

### Phagocytic protists facilitate virus infections in larval zebrafish

Because protists and other microfauna are key to nutrient cycling in aquatic ecosystems^3,9^, we tested whether they might also play a role in the transmission of aquatic viruses. We selected a panel of common freshwater eukaryotes from Carolina Biological Supply Company and prioritized species consumed by larval zebrafish and/or ones identified in genomic data as potentially being associated with zebrafish (**Supplemental figure 1**). Each sample was incubated with zebrafish picornavirus 1 (ZfPV-1; ∼12.5×10^6^ virus genomes)^22,23^, washed to remove free virus, and added to wells of larval zebrafish at 6 days post fertilization (dpf) to screen for the organism’s ability to facilitate virus infections in a vertebrate host (**Figure 1A**).

**Figure 1.**
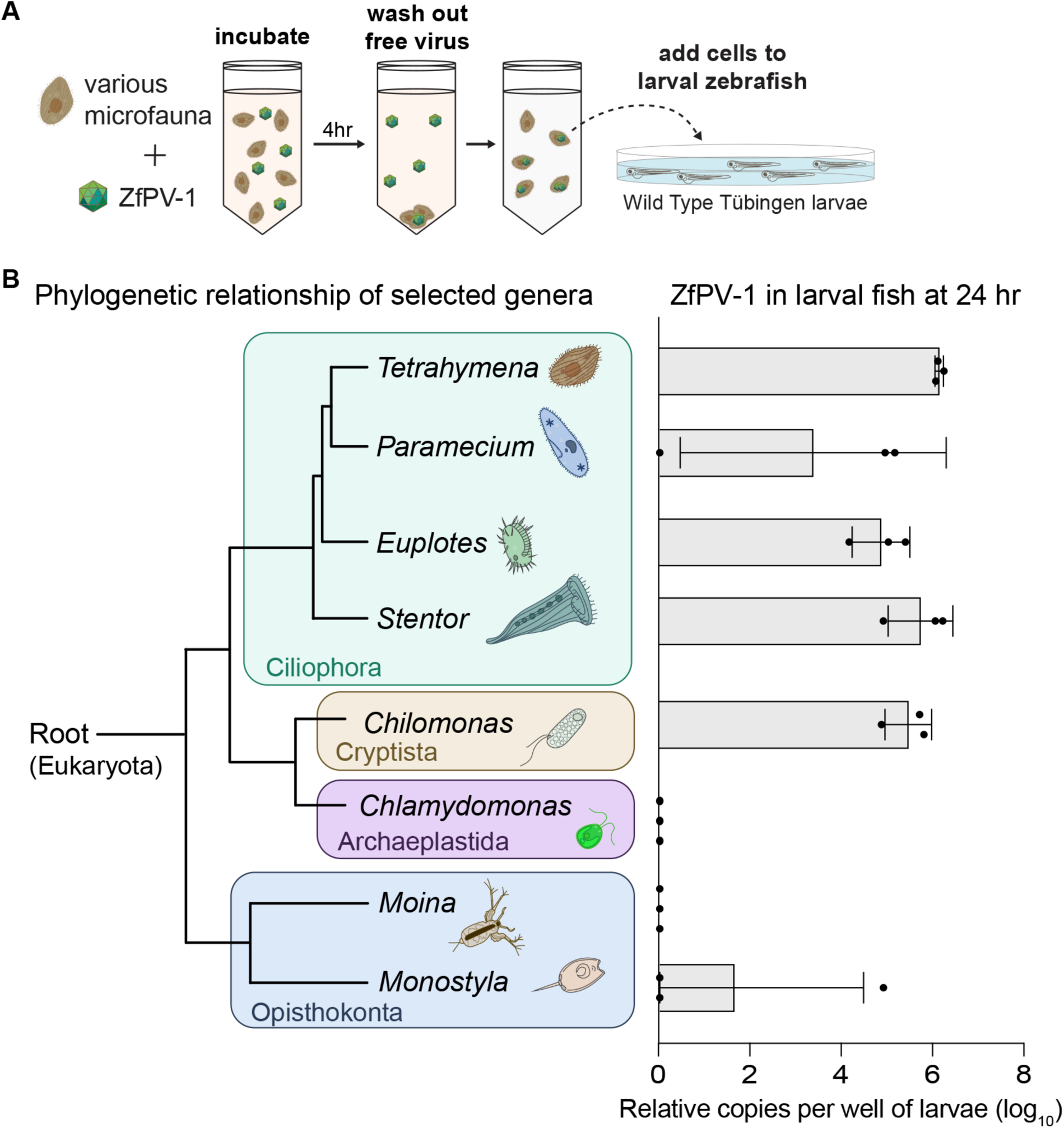
Phagocytic protists acquire ZfPV-1 from water and facilitate transmission to larval zebrafish. **(A)** Experimental workflow for screening a sample of aquatic microfauna for the ability to deliver infectious ZfPV-1 particles to larval zebrafish. Each species was independently incubated with ZfPV-1, washed to remove free virus, and added to wells containing wild type larval zebrafish. **(B)** ZfPV-1 genome copies detected in zebrafish 24 h after exposure to each microeukaryote species. Viral genomes were quantified by qPCR using a standard curve and normalized to the reference gene *eef1a1l1*. Each point represents one pooled sample of 15 larvae. The accompanying phylogenetic tree illustrates the evolutionary relationships among the tested species and was generated based on published phylogenetic data with the acknowledgement that placement of cryptomonads is still controversial.

Each of the four ciliates (*Tetrahymena pyriformis*, *Paramecium tetraurelia, Stentor* sp., and *Euplotes* sp.) and one cryptomonad flagellate (*Chilomonas* sp.) delivered infectious virus, as judged by detection of ZfPV-1 genomes by quantitative PCR (qPCR) in larval fish after 24 hours (**Figure 1B**). In contrast, no ZfPV-1 reads were detected in zebrafish larvae from wells containing the non-phagocytic organisms: *Moina* sp., *Chlamydomonas* sp., and *Monostyla* sp., except for one outlier replicate. While the phagocytic species are only distantly related, these single-celled eukaryotes all consume bacteria, detritus, and other nutrients, suggesting that grazing behavior might play an important role in aquatic virus accumulation. These results suggest an ecologically relevant phenomenon of aquatic virus collection and delivery between phagocytic single-celled eukaryotes and virus hosts.

### The model ciliate *Tetrahymena* accelerates infections and influences antiviral responses in larval fish

To build an experimentally accessible system for studying how single-celled eukaryotes influence aquatic infection dynamics in fish populations, we focused on the model ciliate, *Tetrahymena thermophila*^24,25^. First, we compared viral loads and antiviral responses in larval zebrafish either immersed in ZfPV-1 (a standard route of infection for fish; 2.5×10^5^ virus genomes per 15 larvae) or mixed with *Tetrahymena* that were pre-incubated with an equal amount of ZfPV-1 and washed to remove free virus (**Figure 2A**). To visually compare antiviral responses, we engineered a transgenic zebrafish line to express green fluorescent protein (GFP) in cells producing the interferon-stimulated gene (ISG) *isg15^ref^*^22^ coupled with mCherry fluorescence in the eye as a stable marker of transgenesis (*Tg(isg15:GFP, cryaa:mCherry*).

**Figure 2.**
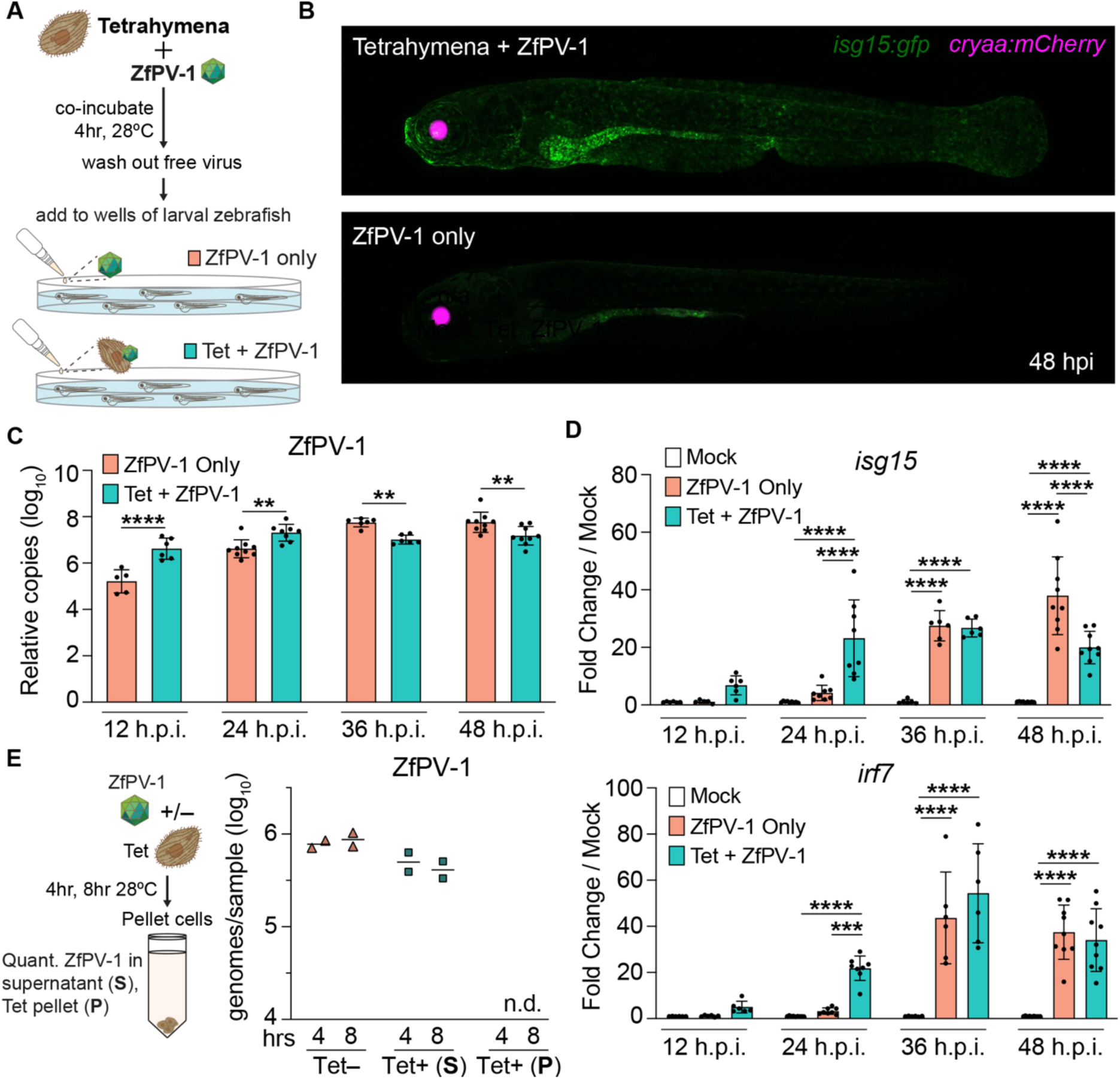
*Tetrahymena*-mediated ZfPV-1 delivery alters infection kinetics and host antiviral responses. **(A)** Experimental workflow to study the ability of *Tetrahymena* to take up and deliver infectious ZfPV-1 particles to larval zebrafish. **(B)** Representative confocal images of *Tg(isg15:GFP, cryaa:mCherry)* larvae (8 dpf) 48 h after exposure to *Tetrahymena* pre-incubated with ZfPV-1 (top) or immersion in ZfPV-1 alone (bottom). **(C)** ZfPV-1 genome copies in pools of 15 larvae over time following infection. Viral genomes were quantified by qPCR using a standard curve and normalized to *eef1a1l1*. Data were analyzed by two-way ANOVA (group × time) followed by Šídák’s multiple-comparisons test within each time point. **(D)** Fold change in interferon-stimulated gene (ISG) expression relative to mock infected fish over time. Data were analyzed by ordinary two-way ANOVA (group x time) with Tukey’s multiple comparisons test comparing groups within each time point. Each point represents one pooled well of 15 individuals from two (12 hpi and 36 hpi) or three (24 hpi and 48 hpi) independent experiments. **(E)** ZfPV-1 genomes were quantified by qPCR from the supernatant and cell pellet after 4 and 8 h of incubation in the presence of absence or *Tetrahymena*. Virus copies normalized to sample volume. Each point represents an independent experiment.

At 48 hours post infection (hpi), zebrafish larvae given virus-incubated *Tetrahymena* exhibited higher levels of GFP expression compared to those immersed in ZfPV-1 alone, with a notably enriched signal in intestines, a primary site of ZfPV-1 replication^22^, as well as the surfaces of the head and trunk (**Figure 2B**). Fish exposed to *Tetrahymena* in the absence of ZfPV-1 resembled uninfected fish, with no detectable GFP. These findings revealed heightened immune responses in larvae exposed to *Tetrahymena* preincubated with ZfPV-1.

To better understand how *Tetrahymena*-virus interactions might impact early infection kinetics, we measured ZfPV-1 genomes in larvae during the initial 48 hours of infection. Compared to ZfPV-1-only fish, *Tetrahymena* delivery of the virus led to significantly more ZfPV-1 in larvae through 24 hpi (**Figure 2C**). This early increase in virus coincides with a drastic increase in ISG expression (**Figure 2D**). Bulk RNA sequencing of larvae at 24 hpi confirmed that *Tetrahymena*-mediated virus infection drives an amplified immune response, rather than a distinct antiviral program, relative to fish infected through immersion in ZfPV-1 alone (**Supplemental Figure 2**). We confirmed these larvae respond to replicating ZfPV-1 because UV-inactivated virus elicited no ISG response in the fish (**Supplemental Figure 3**).

Virus levels and ISG expression in the ZfPV-1-only fish appear to catch up around 36 hpi, suggesting ciliate-based virus delivery accelerates the course of the infection. It is worth noting that the apparent discrepancy between GFP expression (**Figure 2B**) and *isg15* transcripts (**Figure 2D**) at 48 hpi is likely due to differing kinetics of transcription and protein accumulation. Collectively these data reveal that virus-ciliate interactions facilitate ZfPV-1 infections in zebrafish and lead to rapid and robust antiviral responses in these fish.

### ZfPV-1 does not replicate in *Tetrahymena*

Picornaviruses typically exhibit a restricted host range due to host cell receptor specificity and replication machinery requirements ^26–28^, making it unlikely that *Tetrahymena* acts as a biological vector to facilitate ZfPV-1 transmission as an intermediary host for virus replication.

Nevertheless, we assayed for ZfPV-1 replication in *Tetrahymena* and found no increases in ZfPV-1 genome copies at relevant time points (**Figure 2E**). In addition, the amount of ZfPV-1 in the *Tetrahymena* pellet remained below the limit of detection, suggesting the cells contain relatively few particles at any given time point. These data distinguish the role for ciliates in aquatic virus transmission from virus vectors and support the idea that many forms of facilitated virus transmission are likely to evolve in various ecosystems.

### Active phagocytosis by *Tetrahymena* amplifies antiviral responses in larval zebrafish

As opportunistic grazers, ciliates continually consume nutrients and other particles through phagocytosis at a mouth like opening called the oral apparatus. To test whether *Tetrahymena* take up virus particles through phagocytosis, we blocked the process with cytochalasin B and colchicine treatment^29,30^, as confirmed by lack of India ink consumption^31^ (**Figure 3A**). Using the experimental workflow shown in **Figure 2A** with a one-hour virus incubation step following *Tetrahymena* drug treatment, we found that fewer groups of larval zebrafish had detectable ZfPV-1 at 24 hpi relative to untreated cells, as indicated by repeated instances where ZfPV-1 was undetected in the pooled larvae (**Figure 3B**). This finding suggests ciliate-virus interactions may be partially mediated by phagocytosis. Strikingly, there was no detectable antiviral response in any fish population receiving phagocytosis-deficient cells, indicating that the ISG response depends on the ability of *Tetrahymena* to take up viruses by phagocytosis (**Figure 3C**), even though most zebrafish groups had ZfPV-1 levels comparable to ones given untreated *Tetrahymena*.

**Figure 3.**
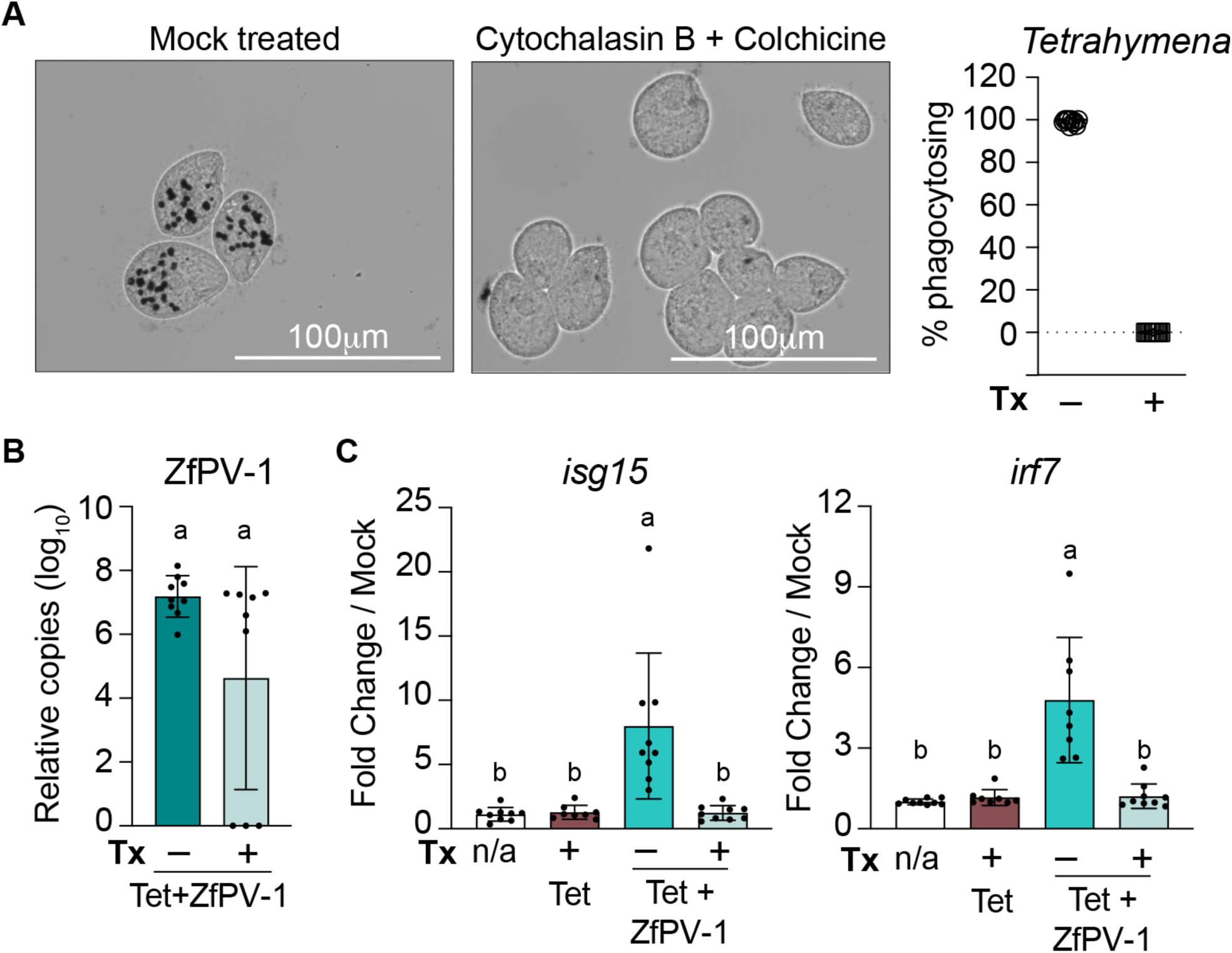
*Tetrahymena* phagocytosis of ZfPV-1 drives zebrafish ISG responses. **(A)** Representative brightfield images of India ink uptake by untreated *Tetrahymena* and cells treated with 12.5μM cytochalasin B and 0.21μM colchicine for 60 min. Phagocytosis was quantified as the percentage of cells containing India ink. Each point represents a blinded quantification from a single image containing 50–100 cells. **(B)** ZfPV-1 genome copies per well of 15 pooled larvae. Viral genomes were quantified by qPCR using a standard curve and normalized to *eef1a1l1*. Values at zero indicate samples that fell below the qPCR limit of detection (approximately 200 genome copies per qPCR reaction). Data were analyzed using Welch’s t test. **(C)** Fold change in ISG expression in larval zebrafish following exposure to untreated or phagocytosis-inhibited *Tetrahymena*. Data were analyzed by ordinary one-way ANOVA with Tukey’s multiple comparisons test. For B-C, each point represents one pooled sample of 15 larvae from three independent experiments.

Next, we tested whether continued association between viruses and live ciliates was required to drive the rapid infection and immune response. After ZfPV-1 incubation, *Tetrahymena* were lysed through repeated freeze-thaw cycles. Compared to larval fish receiving intact *Tetrahymena*-ZfPV-1, we observed a sharp reduction in ISG expression while virus levels remained largely consistent across experimental groups (**Figure 4A and 4B**).

**Figure 4.**
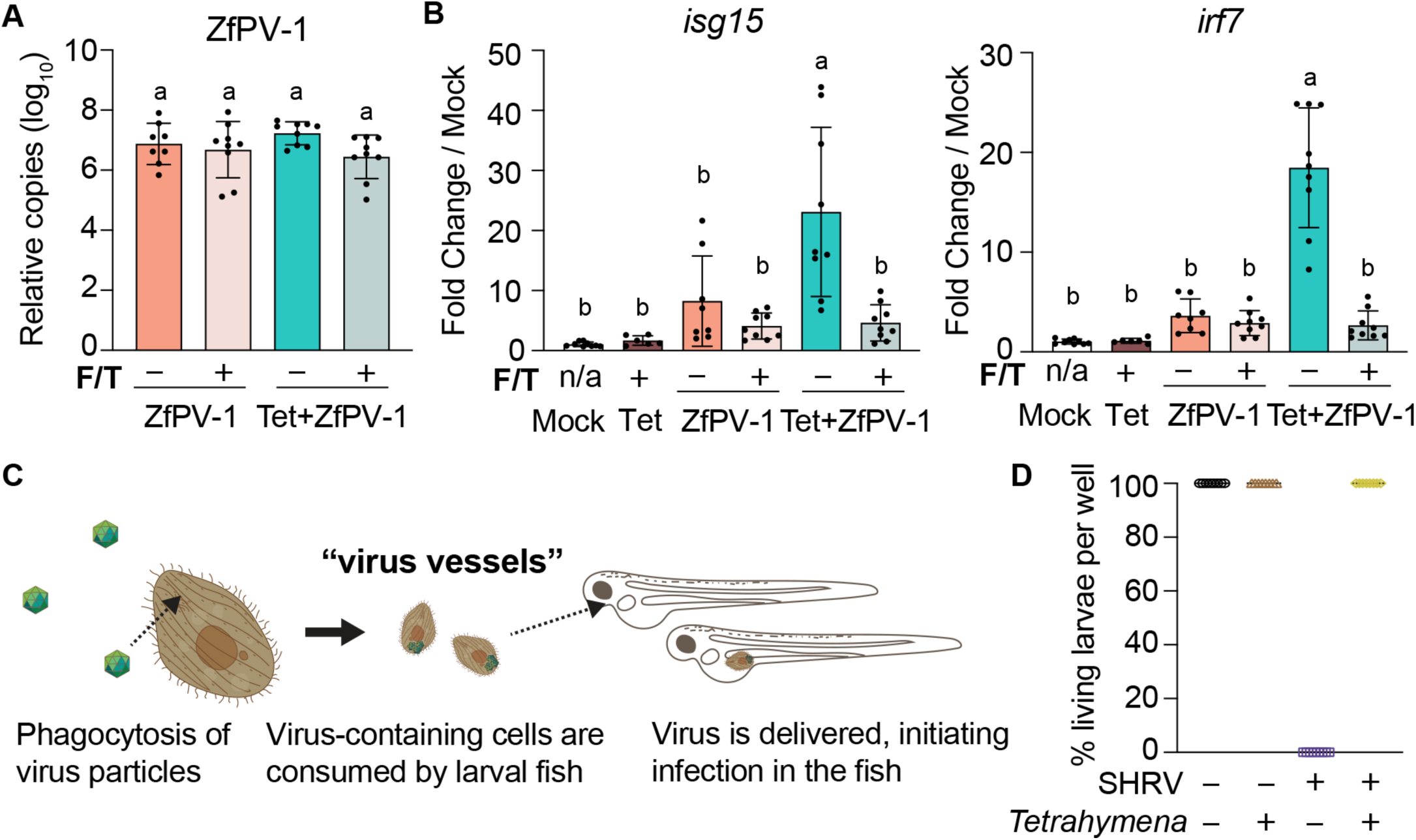
*Tetrahymena* function as vessels for ZfPV-1 transmission. **(A)** ZfPV-1 genome copies detected in pools of 15 larvae following exposure to intact or freeze-thaw (F/T)-lysed *Tetrahymena*. Viral genomes were quantified by qPCR using a standard curve and normalized to *eef1a1l1*. F/T indicates cell lysis by three successive freeze-thaw cycles. **(B)** Fold change in ISG expression relative to mock-infected larvae. Data from panels A and B were analyzed by ordinary one-way ANOVA with Tukey’s multiple comparisons test. Each point represents one pooled sample of 15 larvae from three independent experiments. **(C)** Proposed model in which phagocytes like *Tetrahymena* act as virus vessels by take acquiring particles through phagocytosis and delivering them to susceptible virus hosts, likely as the host consumes virus-laden cells. **(D)** Survival of larval zebrafish 48 h after exposure to snakehead rhabdovirus (SHRV) alone or SHRV-associated *Tetrahymena*. Each point represents one pool of 15 larvae.

The variation observed for immune stimulation versus virus transmission in both sets of experiments suggests additional means of virus uptake by *Tetrahymena*, possibly by clathrin-mediated endocytosis, which is distinct from phagocytosis at the oral apparatus^14,32^. Taking these results together, we propose the term virus vessels for phagocytic protists like *Tetrahymena* that act to promote infections, but do not serve as intermediate hosts for virus replication, as in the case of virus vectors (**Figure 4C**).

### Distinct outcomes of ciliate-mediated transmission for different viruses

To test if *Tetrahymena* act as vessels for the transmission of other fish-infecting viruses, we incubated cells with snakehead rhabdovirus (SHRV). Unlike ZfPV-1, SHRV is highly pathogenic to many fish species, including zebrafish^33,34^, providing a conspicuous phenotype of infection. Using the experimental setup described in **Figure2A,** we observed 100% mortality in larvae immersed in 250 50% tissue culture infective doses (TCID_50_)/mL of SHRV by 18 hpi (**Figure 4D**). In sharp contrast, none of the larvae given *Tetrahymena* pre-incubated with the same dose of SHRV showed any signs of pathology even two days after infection, and none of the larvae yielded detectable SHRV by qPCR. These experiments suggest that SHRV is degraded by phagocytosis, like other viruses^10–14^, and revealed that ciliate-facilitated virus transmission is not ubiquitous across RNA fish viruses. In contrast, another type of aquatic virus, chum salmon reovirus, increases in ability to replicate after phagocytosis by *Tetrahymena* ^35^, hinting at the possibility that ZfPV-1 and other viruses coevolve to benefit from associations with virus vessels.

### *Tetrahymena* sustains chains of virus transmission across zebrafish populations

To test if ciliate-virus interactions by vessels like *Tetrahymena* impact virus transmission across host populations over time, we set up an experimental system to model virus transmission between larval fish. To begin, a group of “virus donor” larvae (*Tg(isg15:GFP, cmlc2:GFP*))^22^ were infected through immersion in ZfPV-1. At 48 hpi, GFP^+^ infected larvae were selected, washed, and added to uninfected “virus recipient” larvae (*Tg(isg15:GFP, cryaa:mCherry*)) in the presence or absence of *Tetrahymena* (**Figure 5A, left of the dashed line**). Heart-specific *cmlc2*-driven expression of GFP versus eye-specific cryaa-driven expression of mCherry allowed for quick, visual scoring of virus donor versus recipient fish.

**Figure 5.**
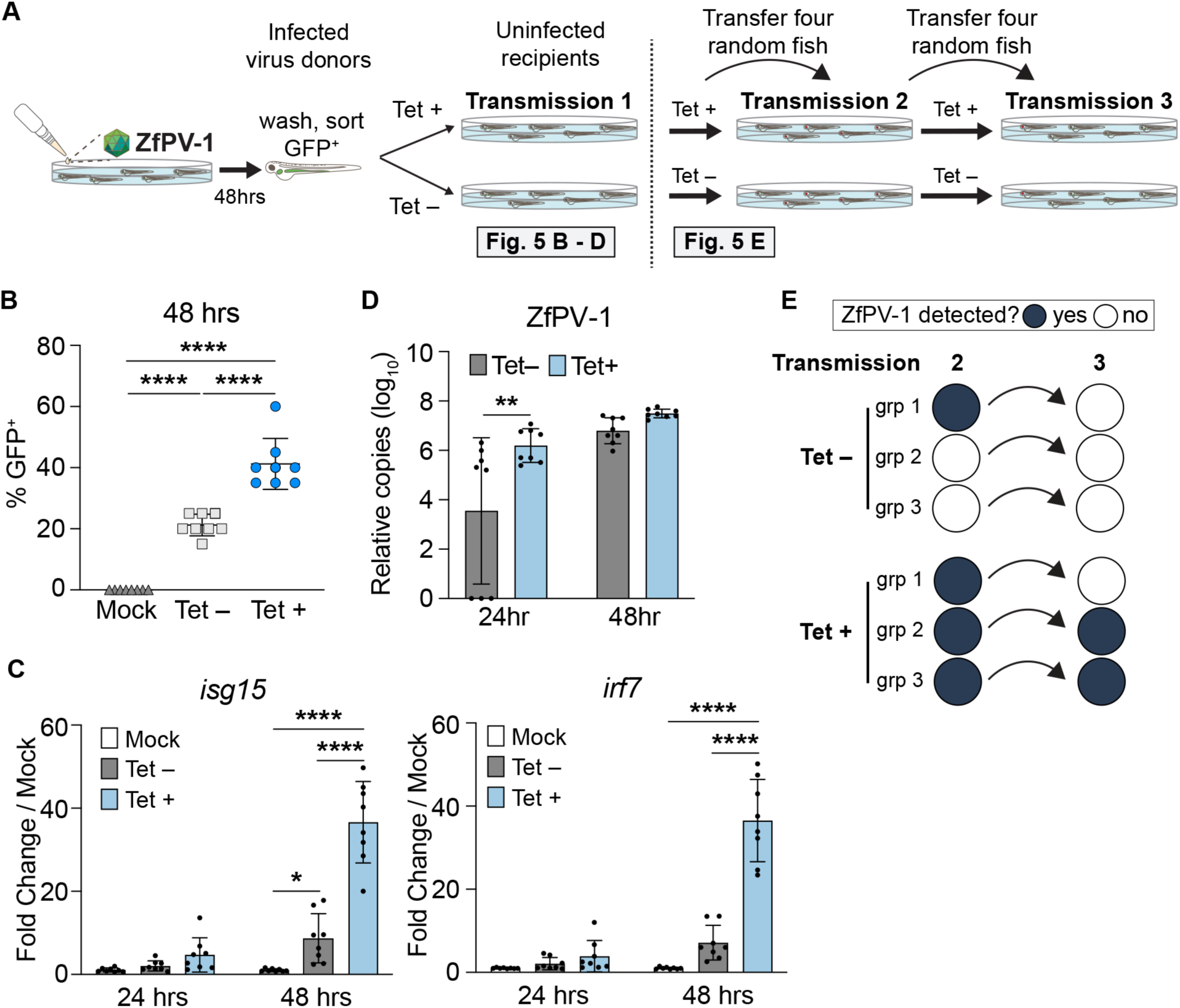
Tetrahymena sustains experimental chains of ZfPV-1 transmission between zebrafish populations. **(A)** Schematic of sequential transmission assays. **Left**: Pools of donor *Tg(isg15:GFP, cmlc2:gfp)* larvae were infected through immersion. At 48 hpi, infected fish were identified by robust GFP expression in the intestine and co-housed with naïve *Tg(isg15:GFP, cryaa:mCherry)* larvae in the presence or absence of *Tetrahymena*. Primary transmission data are shown in panels B and C. **Right**: For sequential transmission events, four recipient larvae were randomly selected after each co-housing period and transferred to a new population of naïve larvae. ZfPV-1 genomes were quantified for each population in the second and third transmissions. **(B)** Percent of GFP-positive recipient larvae at 48 hours post co-housing with ZfPV-1 infected or mock infected virus donor fish in the presence or absence of *Tetrahymena*. Data were analyzed by ordinary one-way ANOVA with Tukey’s multiple comparisons test. **(C)** Fold change in ISG expression relative to mock per pool of larvae. Data were analyzed by ordinary two-way ANOVA (group x time) with Tukey’s multiple comparisons test comparing groups within each time point. **(D)** ZfPV-1 copies per well of 20 pooled larvae. Data were analyzed by ordinary two-way ANOVA (group x time) followed by Šídák’s multiple comparisons test within each time point. For C-D, each dot represents one pooled well of 20 individuals across two independent experiments. **(E)** Depiction of ZfPV-1 transmission between three independent groups of larval zebrafish in the absence (top) or presence (bottom) of *Tetrahymena*. Filled circles represent wells where ZfPV-1 genomes were detected by qPCR in the fish pooled from that well. Quantification is provided in Supplemental Figure 4.

Remarkably, after 48 hours of cohousing, there were significantly more GFP^+^ recipient fish in wells containing *Tetrahymena* than wells without (**Figure 5B**), indicating the presence of ciliates increased virus transmission between fish. Larvae co-housed in the absence of *Tetrahymena* exhibited GFP expression in intestines, while those infected in the presence of *Tetrahymena* had a more widely distributed GFP signal akin to fish shown in Figure 2B. Corroborating the GFP data, there was a drastic increase in the expression of *isg15* and *irf7* transcripts (**Figure 5C**) in these larvae at 48 hours of infection. Although there was little visible GFP at 24 hours post cohousing, qPCR revealed ZfPV-1 genome copies in both populations of recipient fish at this earlier time point, with a higher viral load in larvae cohoused in the presence of *Tetrahymena* (**Figure 5D**). These results show that the simple presence of *Tetrahymena* in infected and uninfected populations of hosts can facilitate virus transmission.

To extend experimental chains of virus transmission over multiple rounds of infection, we selected four random larvae from each “virus recipient” group to become potential virus donors in a subsequent encounter with uninfected fish, repeating this process for a total of two additional rounds of transmission in the presence or absence of *Tetrahymena* (**Figure 5A, right of the dashed line**). The presence of ciliates had a clear impact on continued transmission. ZfPV-1 genome levels dropped below the limit of qPCR detection in two of the three experimental groups in the absence of *Tetrahymena* by the second transmission event, with virus completely lost by the third round of transmission (**Figure 5E** and **Supplementary Figure 4 A**). However, in the presence of *Tetrahymena*, ZfPV-1 was detected in all zebrafish groups in the second round of transmission and in two of the three groups by the third round (**Figure 5E** and **Supplementary Figure 4A**). Quantification of GFP expression and ISG transcripts in recipient fish reflected the rapid loss of infectious virus in the absence of the ciliates (**Supplementary Figure 4B, C**). Together these results reveal the pivotal role phagocytic cells like *Tetrahymena* can play in sustaining chains of virus transmission across vertebrate host populations.

## Discussion

Pathways of virus transmission vary widely. Establishing experimental systems to study modes of transmission will improve our understanding of how viruses sustain propagation through host populations—a fundamental need for mitigating the spread of infectious diseases. Here we model virus transmission among fish populations, where particles encounter numerous phagocytic, non-host organisms that may directly aid or inhibit virus propagation.

These phagocytic cells, including ciliates, provide potential means of facilitating virus transmission by concentrating and delivering particles. However, phagocytosis initiates enzymatic digestion, and previous work instead points to ciliates being barriers to virus transmission with a suite of proteolytic enzymes poised to derive energy from collected particles, including viruses^10–14^. In this study, the complete loss of viral pathogenesis of snakehead rhabdovirus after mixing with *Tetrahymena* (Figure 4D) is consistent with virus particles being consumed and degraded.

In contrast, some viruses seem adapted to the onslaught of digestive enzymes after phagocytosis, including chum salmon reovirus, which appears to be processed into a more infectious form after association with *Tetrahymena*^35^. For zebrafish picornavirus 1 (ZfPV-1), the outcome for ciliate-mediated transmission appears more complicated. *Tetrahymena*-based virus delivery initiates fast and robust immune responses in zebrafish, coupled with modest increases in ZfPV-1 genomes. Most striking from this study, however, was an experimental dependence on *Tetrahymena* for sustaining chains of ZfPV-1 transmission (Figure 5). This finding suggests that ciliates sustain certain virus-host relationships found in natural settings.

The distinct infection outcomes from non-host encounters reported here and elsewhere may be driven by core cell biological features of ciliates, including differentiated pathways of membrane trafficking. While particles taken up through phagocytosis might be largely degraded into immunostimulatory cues, the independent process of clathrin-mediated endocytosis might provide a route less vulnerable to immediate proteolysis^14,32,36^. Studying if and how virus populations adapt to navigate ciliate cell biology will likely be a fruitful direction for future work.

The discovery of ciliates as virus vessels for ZfPV-1, an endemic virus of zebrafish, points to a vast middle ground between direct host-to-host virus transmission and replication in intermediary virus vectors. Arthropod-facilitated transmission of various plant viruses shows this continuum also exists in terrestrial systems^37–39^, suggesting non-host mediated transmission as a wide-spread phenomenon that warrants further research. Ultimately, by layering together research organisms with scores of experimental tools, we build a pathway beyond traditional reductionist approaches to better understand the process of virus transmission, which holds great promise for future multidisciplinary studies of infection biology.

## Methods

### Zebrafish care and larvae generation

All zebrafish (*Danio rerio*) used in this study were kept and maintained in accordance with the approved Institutional Animal Care and Use Committee (IACUC) Protocol #00001439. All adult and larval zebrafish were kept on a 14-10 hour light-dark cycle at 28°C. Wild type Tübingen (Tü) fish were acquired from the University of Utah’s Centralized Zebrafish Animal Resource. *Tg(isg15:GFP, cmlc2:GFP)* fish were generated previously^22^. *Tg(isg15:GFP, cryaa:mCherry)* fish were similarly generated using the Tol2kit^40^ and backcrossed to wild type Tü a minimum of three times before being used in any experiments. All zebrafish lines were maintained through intercrossed matings and/or outcrossing to wild type Tü fish.

Embryos for each experiment were generated through in-cross matings of indicated zebrafish lines. All embryos were surface sanitized 24-30 hours post-fertilization (hpf) in a 30 parts per million sodium hypochlorite solution, made fresh from a concentrate containing 10-15% available chlorine (Milipore Sigma), following a standard sanitization protocol. Specifically, each clutch was immersed for five minutes in the sodium hypochlorite solution followed by a five-minute immersion in deionized water, repeated twice with fresh solutions. The embryos were then washed by two subsequent five-minute immersions in E3 media (5 mM NaCl, 0.17 mM KCl, 0.33 mM CaCl2, 0.33 mM MgSO4). Sterilized embryos were transferred to sterile Petri dishes and dechorionated using a 2mg/mL Pronase solution in E3 media (Sigma-Aldrich). Dechorionated embryos were reared on a 14-10 hour light-dark cycle at 28°C in 10 cm dishes containing 60 mL E3.

One day prior to infection, 5-day post-fertilization (dpf) larvae were lightly anesthetized with buffered tricaine mesylate (Syncaine/Tricaine/MS-222, Syndel) at a final concentration of 200 mg/mL and sorted into 6 well non-treated plates containing 5mL of fresh E3 media at a density of 15 larvae per well, unless otherwise noted. Equal numbers of larvae from multiple clutches were added to each well to avoid potential clutch-to-clutch variation in infection. All larvae were fed live rotifers starting at 6 dpf.

### Zebrafish picornavirus 1 (ZfPV-1) stock generation

Zebrafish picornavirus 1 (ZfPV-1) was harvested from intestines of adult zebrafish as previously described^22^ with minor changes. Adult zebrafish (1-2 years old) were fasted one day prior to harvest to reduce food contents in the intestine. Fish were euthanized through ice water immersion and secondary decapitation in accordance with IACUC Protocol #00001439. Six to seven intestines were pooled per 3 mL of E3 media, homogenized with a THb Handheld Tissue Homogenizer, and subject to three rapid freeze/thaw cycles. Homogenates were then pooled and spun at 10,000 x *g* for 10 minutes to clear large debris. Smaller debris was removed by filtration through a 0.45 mm Steriflip^®^ Vacuum Tube Top Filter (Milipore Sigma). Cleared lysate was aliquot and stored at –80°C. ZfPV-1 genome copies were quantified through quantitative PCR using a standard curve.

### Snakehead rhabdovirus stock generation and quantification

Snakehead rhabdovirus stock was acquired from ATCC (VR-1386). ZF4 cells (CRL-2050, ATCC) were grown to confluence in T-175 tissue culture treated flasks (Thermo Scientific, 159910) in DMEM:F12 (Gibco, 11320082) supplemented with 10% heat inactivated fetal bovine serum (Sigma Aldrich, F0926-100ML) at 28.5°C with 5% CO_2_. The media was aspirated and adherent cells were washed once with phosphate buffered saline (diluted from 10X PBS pH 7.4, Gibco, 70011-044). SHRV stock was mixed at 50μL/mL in DMEM:F12, and a total of 15mL was added to each T-175 flask. Flasks were rocked every 15 min for one hour following infection, then left static overnight at 28.5°C. Upon observation of >90% cytopathic effect, media and cells were harvested using a cell scraper. Virus-free controls were subjected to one freeze/thaw at –80°C/+37°C. Solutions were centrifuged at 500 x g for 5 min at 4°C, and supernatants were collected, aliquoted, and stored at –80°C. Virus stocks were titered using a TCID_50_ endpoint titration assay (Spearman–Kärber method) in ZF4 cells^41^.

### Maintenance of ciliates and other microfauna

*Tetrahymena pyriformis*, *Paramecium tetraurelia, Stentor* sp., *Euplotes* sp., *Chilomonas* sp., *Moina* sp., *Monostyla* sp., and *Chlamydomonas* sp. were purchased from Carolina Biological Supply. Upon arrival, cultures were examined for viability and used within two hours of receipt. All experiments except the initial microfauna screening used *Tetrahymena thermophila* (strain SB210, RRID: TSC_SD00703), which was purchased from the Tetrahymena Stock Center and maintained in the laboratory in Modified Neff Medium (0.25% proteose peptone, 0.25% yeast extract, 0.5% glucose, 33.3 mM FeCl3) at room temperature through routine serial passaging.

### *Tetrahymena* mediated virus transmission assays

*Tetrahymena* were cultured to mid-log growth phase by shaking at 125 rpm overnight at 30°C in non-baffled flasks. The following morning, cells were harvested by centrifugation (5 min, 300 x *g*), resuspended in fresh Neff Medium, and adjusted to a density of 3×10^4^ cells per mL. *Tetrahymena* were incubated with ZfPV-1 (2.5×10 genomes per 1×10 cells) or mock-treated in Modified Neff medium for 4 h at 28°C unless otherwise indicated. Cells were then pelleted by centrifugation (5 min, 300 x *g*), washed three times in E3 media to remove free virus, and 1×10^4^ cells were added to each well containing 15 larval zebrafish. For ZfPV-1 only conditions, virus (2.5×10^5^ ZfPV-1 genomes per pool of 15 fish) was added directly to the larvae. Equal volumes of E3 were maintained across all zebrafish conditions. In experiments extending beyond 24 hours post infection (hpi), larvae received twice daily rotifer feedings and media changes.

At the indicated harvest time points, larvae were euthanized by an overdose of >300 mg/L Tricaine and secondary ice bath immersion and washed three times in MilliQ water to remove ciliates and residual free virus. Each pool of 15 larvae was homogenized together in TRIzol (Invitrogen Life Technologies, 15596018).

### Modifications to *Tetrahymena*-mediated virus transmission assays

#### Microfauna screen

5 mL cultures of *Tetrahymena pyriformis*, *Paramecium tetraurelia, Stentor* sp., *Euplotes* sp., *Chilomonas* sp., *Moina* sp., *Monostyla* sp., and *Chlamydomonas* sp. were each incubated with 12.5×10^6^ ZfPV-1 genome copies for 4 hours at 23°C. All cultured organisms were washed twice in E3 media to remove free virus and resuspended in a final volume of 1mL E3 per pool of larval zebrafish. *Euplotes* sp., *Chilomonas* sp., *Tetrahymena pyriformis*, and *Paramecium tetraurelia*, were pelleted by centrifugation (5 min, 300 x *g*). *Chlamydomonas* sp. was pelleted by centrifugation (3 min, 1500 x *g*). *Monostyla* sp. was pelleted by centrifugation (90 seconds, 100 x *g*). *Monostyla* sp. was gently washed over a 40 μm cell strainer. *Stentor* sp. was washed by allowing the cells to settle and removing the supernatant.

#### ZfPV-1 inactivation by UV irradiation

ZfPV-1 was inactivated by 3 1-min exposures to ultraviolet (UV) light using a Stratalinker 1800 UV Crosslinker (Stratagene, 400071) operated in Time Mode. UV-treated virus was used immediately in downstream assays. Loss of infectivity was verified by a lack of detectible ZfPV-1 genome copies and ISG response in larval zebrafish.

#### Disruption of phagocytosis in Tetrahymena with Colchicine and Cytochalasin B

Colchicine (Millipore, C9754-500MG) was dissolved in 10 mM HEPES buffer, pH 7.0 to 21μM and used at a final concentration of 0.21μM. Cytochalasin B (Millipore, C6762-5MG) was dissolved in DMSO to 25μM and used at a final concentration of 12.5μM. *Tetrahymena* cells were pre-incubated with both drugs for 20 minutes at 28°C before the addition of ZfPV-1 or India ink. Because prolonged drug exposure reduced *Tetrahymena* viability, incubations with ZfPV-1 or India ink were limited to 1 hr. To quantify phagocytosis, cells incubated with India ink were fixed with a 5% glutaraldehyde solution and imaged on an AMG EVOS Digital Inverted Microscope. Samples were blinded prior to analysis, and phagocytosis was assessed as the percent of cells containing India ink-filled food vacuoles.

#### Tetrahymena lysis by successive freeze-thaws

Following the final *Tetrahymena* wash step and resuspension in E3 medium, each sample was divided equally into two aliquots. One aliquot served as the intact control, while the other was subject to three rapid freeze-thaw cycles using liquid nitrogen and a room-temperature water bath. *Tetrahymena*-only and ZfPV-1-only control samples underwent identical processing.

#### Snakehead rhabdovirus infections

The *Tetrahymena*-mediated virus exposure assay described above was performed using snakehead rhabdovirus (SHRV) in place of ZfPV-1. Pools of 20 larval zebrafish were exposed to either 250 TCID_50_ of SHRV or *Tetrahymena* pre-incubated with an equivalent amount of virus in 3 mL of E3 medium. Virus-free cell culture supernatant served as the negative control. Larvae were monitored for morbidity and mortality throughout the experiment, and all surviving fish were harvested at 48 hpi.

#### RNA extraction and RT-qPCR

RNA was extracted following the TRIzol Reagent User Guide (Pub. No. MAN0001271), followed by DNase treatment and cDNA synthesis using the Maxima First Strand cDNA Synthesis Kit (Thermo Scientific, FERK1672). qPCR was performed in 10 uL reactions using the Applied Biosystems™ Power SYBR™ Green Master Mix (ThermoFisher, 4368708) and the manufactures recommended standard cycling conditions. Reactions were run on an Applied Biosystems™QuantStudio™ 5 Real-Time PCR System (ThermoFisher, A34322). Absolute ZfPV-1 genome abundance was determined by comparison to a plasmid standard curve and then normalized to *eef1a1l1*. Relative ISG expression was calculated against mock infected fish using the 2^−ΔΔCt^ method with *eef1a1l1* as the endogenous control. Primer sequences are provided below:

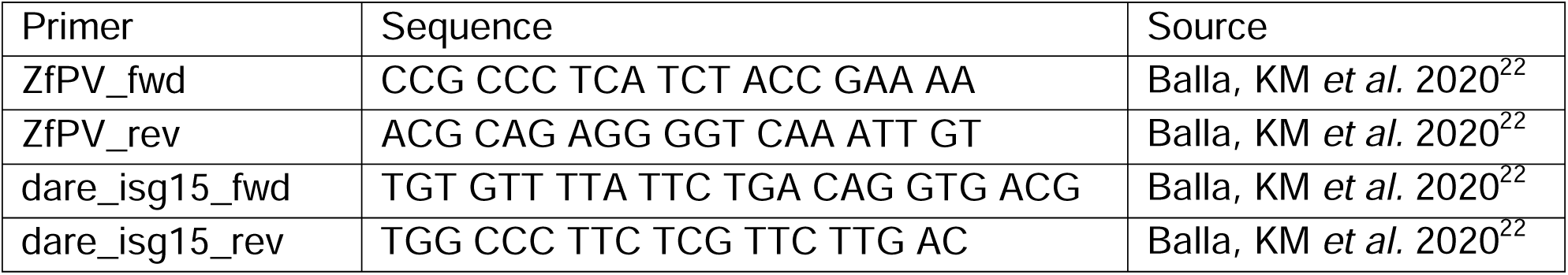
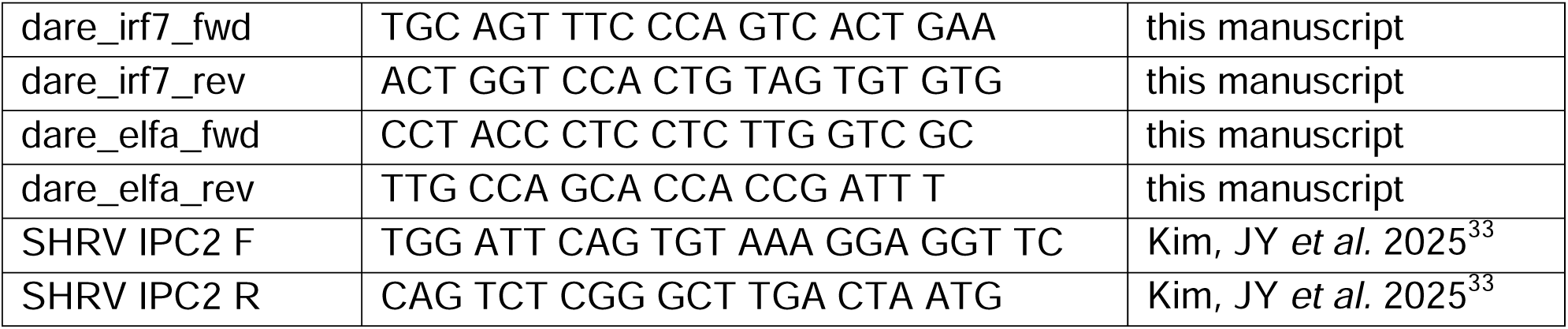

### Confocal imaging

At 48 hpi, larvae were anesthetized with buffered tricaine mesylate (Syncaine/Tricaine/MS-222, Syndel) at a final concentration of 200 mg/mL and mounted in 1% low melt agarose (Lonza). Individual fish were imaged with a Zeiss LSM 980 confocal microscope (Zeiss) with an NA 0.45/10x objective using the 488 nm and 561 nm laser lines. Images were acquired using a 25 µm pinhole size at a resolution of 0.72 pixels per µm. Z-stacks were imaged with a step size of 2.02 µm. Images are maximum intensity projections. Tiling was used to obtain images of whole animals. Tiles were stitched using ZEN blue software (Zeiss) and cropped and rotated in FIJI (ImageJ). No brightness or color adjustments were made. All images were collected on the same day using the same laser power, gain, and exposure.

### RNA sequencing and Differential Gene Expression Analysis

Total RNA was isolated from pooled larval zebrafish 24 hours after infection using the *Tetrahymena*-mediated virus exposure assay described above and extracted as described in *RNA extraction and RT-qPCR*. A ScreenTape Assay (Agilent Technologies) to assess RNA quality gave RIN values ≥9.2 for all samples. Non-stranded RNA-seq libraries were prepared by GENEWIZ (Azenta Life Sciences) using oligo(dT)-based poly(A) enrichment followed by random-primed cDNA synthesis. Libraries were sequenced on an Illumina NovaSeq X Plus platform to generate 2 x 150 bp paired-end reads to a depth of approximately 30 million read pairs per sample. Raw read data are available through the National Center for Biotechnology Informatics (NCBI) Sequence Read Archive (SRA) under BioProject accession number PRJNA1458218.

Adapter sequences and low-quality bases were removed from raw sequencing reads using TrimGalore v0.6.7 (which uses Cutadapt v1.18 and FastQC v0.11.9). Filtered reads were aligned to the *Danio rerio* GRCz11 reference genome (RefSeq accession GCF_000002035.6) and corresponding RefSeq gene annotation using STAR v2.7.8a. STAR’s --quantMode GeneCounts was used to generate gene-level read counts. Gene-level read counts with fewer than 10 total counts across all samples were excluded prior to differential expression analysis. Differential gene expression analysis was performed in R v4.3.3 with DESeq2 v1.42.1. Genes with a false discovery rate (Benjamini–Hochberg adjusted *P* value) < 0.05 and an absolute log2 fold change >1 were considered differentially expressed.

### Sequential Transmission of ZfPV-1

For the initial infections, 2.5×10^5^ ZfPV-1 genomes were added wells of 20 *Tg(isg15:GFP, cmlc2:GFP*) larvae. At 48 hpi, robustly infected larvae expressed high levels of GFP in the intestine^22^. To select these fish as virus donors for transmission, all larvae were lightly anesthetized with buffered tricaine mesylate (Syncaine/Tricaine/MS-222, Syndel, 200 mg/mL) and visualized under a fluorescent stereo microscope. GFP^high^ donor larvae were removed, gently rinsed in fresh E3 media, and three individuals were transferred to each dish containing 20 uninfected recipient larvae in the presence or absence of *Tetrahymena* (1×10^4^ cells per condition).

### Quantification of GFP expression in primary transmission

At 24 and 48 hours post addition of virus donor larvae, half of the recipient population was harvested as described above with the modification that prior to euthanasia, fish were first lightly anesthetized and visualized under a fluorescent stereo microscope to calculate the number of individuals in each pool that were GFP^+^.

### Blinded serial transmission assay

Each transmission event consisted of 24 h of co-housing between donor and recipient larvae in the presence or absence of *Tetrahymena*. Following each co-housing period, all larvae were lightly anesthetized, and donor fish were removed from the population based on transgenic marker expression (either *Tg(cmlc2:GFP) or Tg(cryaa:mCherry)*). Four recipient larvae were picked at random and transferred to a new population of naive 6 dpf larvae expressing the alternate transgenic marker. This blinded procedure was carried out for a total of three transmission events across three independent populations. Following the third transmission event, the remaining recipient larvae were harvested 48 h after the start of co-housing for qPCR analysis of ZfPV-1 genome copies and ISG expression. At 48 hours of co-housing for the fourth transmission, donor fish were removed, and the recipient larvae were harvested for qPCR analysis of ZfPV-1 genome copies and ISGs.

### Quantification and Statistical Significance

Details regarding quantification and statistical analyses can be found within the corresponding figure legends and text. Data are presented as mean ± SD. Differences were considered statistically significant at *P* < 0.05.

## Data availability

The RNA-sequencing dataset generated for this manuscript s publicly available at the NCBI Sequence Read Archive, Bio-Project accession number PRJNA1458218.

## Acknowledgements

We thank the Tetrahymena Stock Center (Washington University) and D. Chalker for advice and support. This work was supported by NIH grant R35GM142950 to JAG, a predoctoral training grant fellowship to CET (T32AI055434), and a Hanna H. Gray Fellowship to KAD. NCE is an investigator of the Howard Hughes Medical Institute.

## Author contributions

Conceptualization, KAD, JAG, NCE; Investigation, KAD, RTT, HJY, CET; Writing, KAD, JAG, NCE; Resources, JAG and NCE, Funding acquisition, JAG and NCE.

---

**Supplemental Figure 1.**
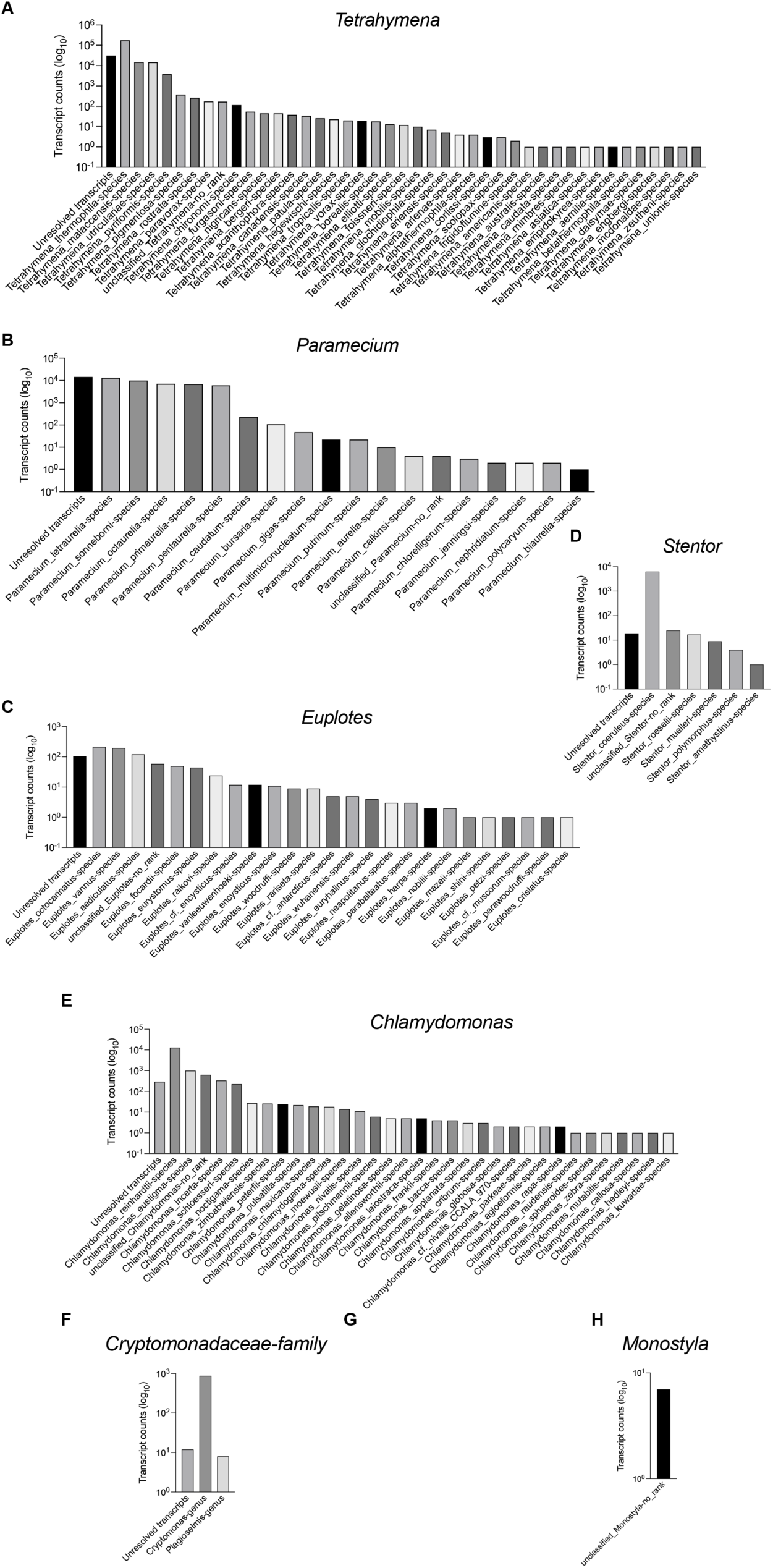
RNAquarium zebrafish data demonstrates wide-spread presence of diverse microeukaryotes. Transcript counts retrieved from the bi[o]hub RNAquarium Phylogenetic Viewer (https://phylotree.rnaquarium.org/ -- accessed 5 August 2026). Organisms were queried at the genus level unless otherwise indicated in the graph title.

**Supplemental Figure 2.**
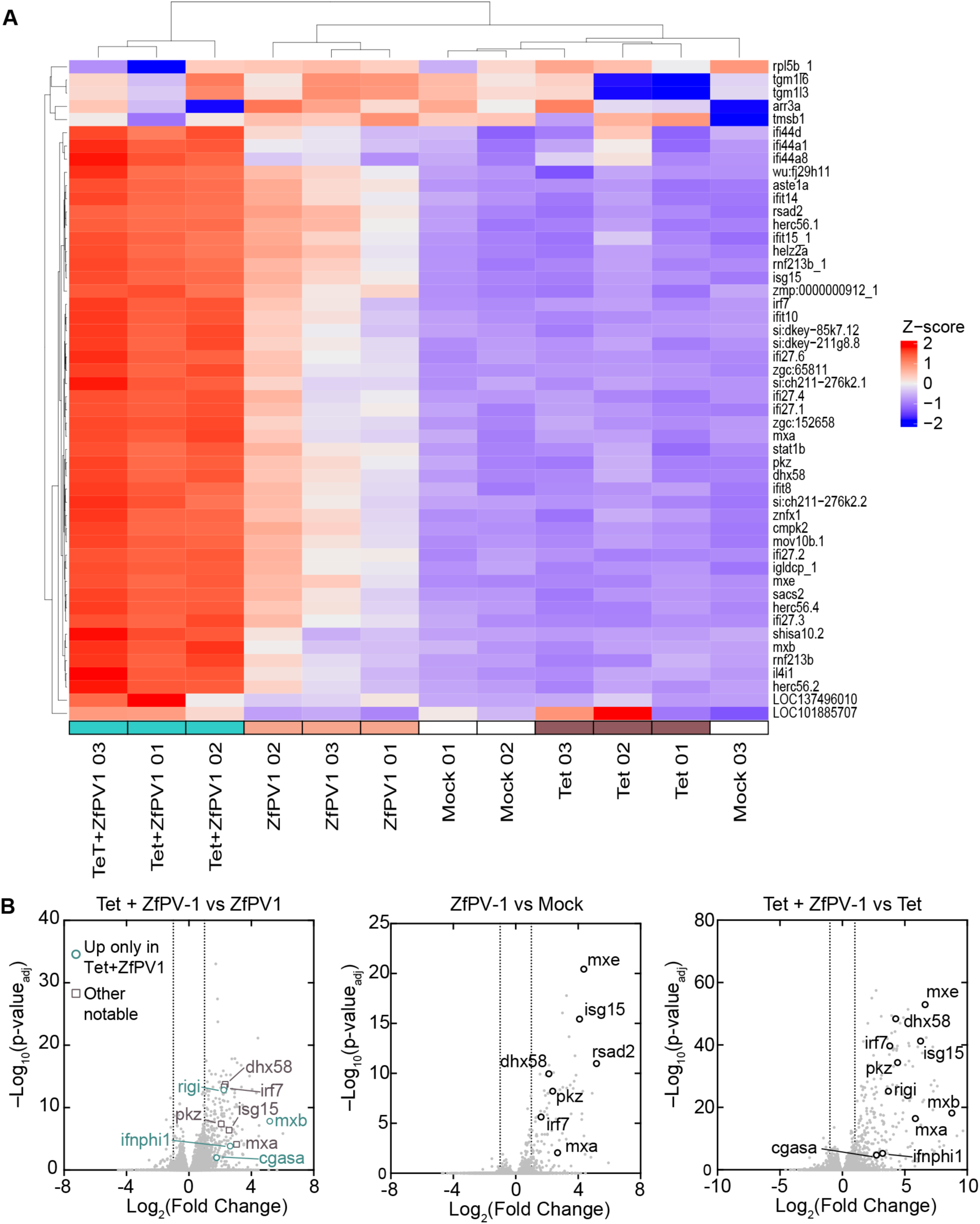
T*e*trahymena-delivered ZfPV-1 drives an amplified antiviral response in larval zebrafish relative to infection with ZfPV-1 alone. (A) Heatmap from RNA sequencing data of the top 50 differentially expressed genes determined by adjusted *P* value (rows) across each sample (columns). Genes and samples are hierarchically clustered. **(B)** Volcano plots of differentially expressed genes between samples indicated in graph titles. Genes with an adjusted *P* value > 0.05 and |log2 fold change| >1 (indicated by the vertical dashed lines) were considered significant. Each dot represents an individual gene, with select known antiviral genes highlighted in each graph.

**Supplemental Figure 3.**
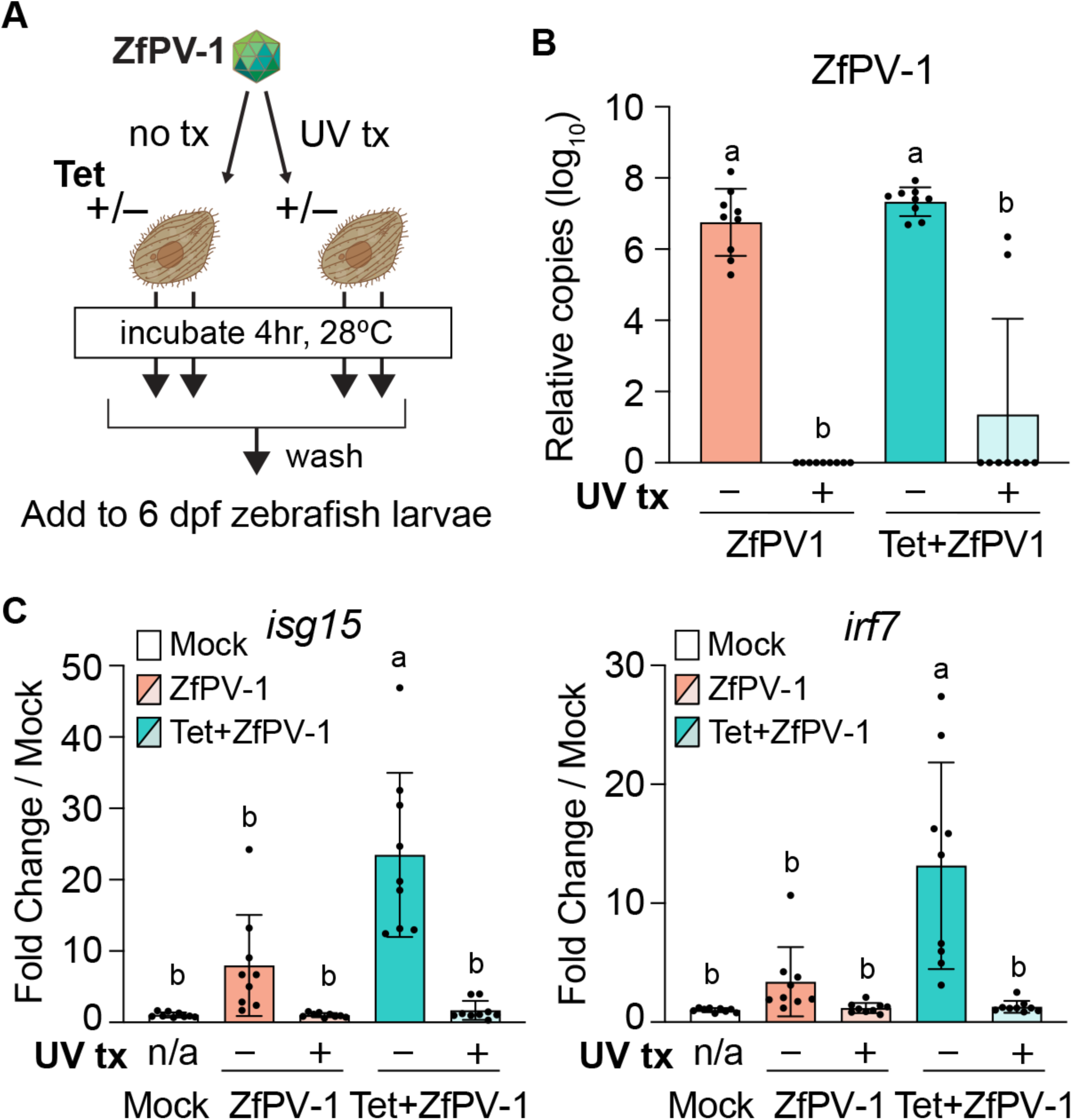
UV treatment of ZfPV-1 eliminates *Tetrahymena*-driven increase in larval ISG response to virus. **(A)** Experimental workflow to determine if increased ISG response in larval zebrafish given *Tetrahymena* pre-incubated with ZfPV-1 is dependent on replication-competent virus. **(B)** ZfPV-1 genome copies. Data points at 0 fell below the qPCR limit of detection (approximately 200 genome copies per qPCR reaction). **(C)** Fold change in ISG expression relative to mock-infected larvae. Data were analyzed by ordinary one-way ANOVA with Tukey’s multiple comparisons test, and each point represents one pool of 15 individuals across three independent experiments.

**Supplemental Figure 4.**
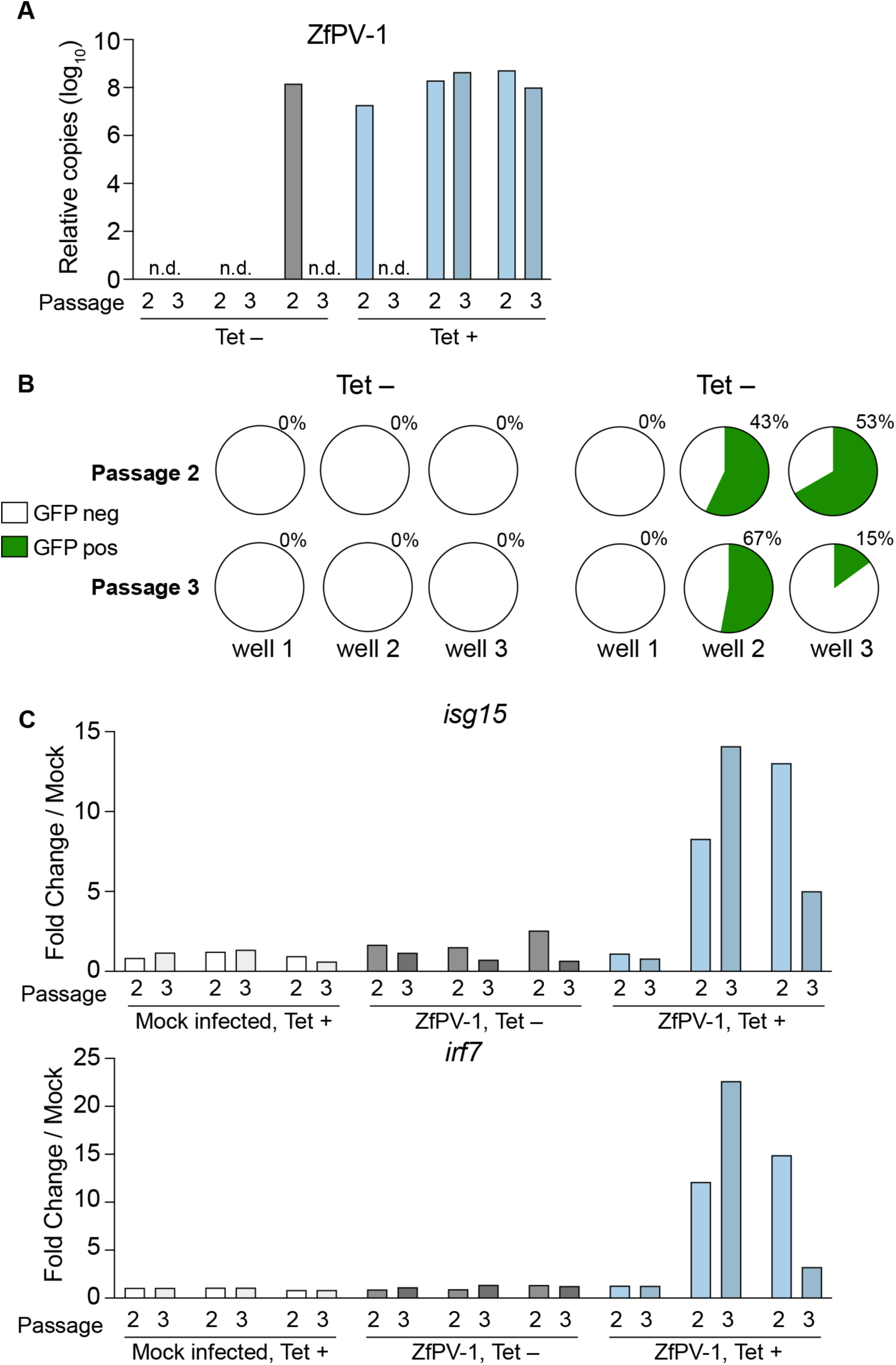
Quantitative analysis of the serial transmission experiments shown in. **Figure 5E. (A)** ZfPV-1 genome copies per pool of 20 larvae, quantified by qPCR using a standard curve and normalized to *eef1a1l1*. Each bar represents one pool of fish. n.d., not detected. **(B)** Percentage of GFP-positive larvae in each pool at each transmission event. Each pie chart represents one pool of larvae, with the percentage of GFP-positive larvae indicated above. **(C)** Fold change in ISG expression relative to mock infected controls at 48 hours after the start of each transmission event. Each bar represents one pool of larvae.

